# Validating Dynamic Body Acceleration metrics as a measure of energy expenditure in a Neotropical primate

**DOI:** 10.1101/2023.06.29.547103

**Authors:** Gabriela C. Rezende, Ariovaldo P. Cruz-Neto, Luca Börger, James Redcliffe, Catherine Hambly, John R. Speakman, Guilherme S. T. Garbino, Alcides Pissinatti, Silvia Bahadian Moreira, Rory Wilson, Laurence Culot

## Abstract

2

Quantifying energy expenditure in free-living primates is fundamentally important yet challenging. Acceleration-based metrics such as Dynamic Body Acceleration (DBA), obtained from accelerometers, are potential proxies for energy expenditure, yet have not been previously validated in primates. Here, we validated DBA in black lion tamarins (*Leontopithecus chrysopygus*) by comparison to doubly labelled water (DLW) in 10 captive tamarins housed at the Rio de Janeiro Primate Centre. Individuals were equipped over 48 hours with a backpack with a tri-axial accelerometer and received an intra-peritoneal DLW injection at the beginning of the experiment, with blood samples taken 1 and 48h later. Daily Energy Expenditure by DLW was 326 (SD=66) kJ/day, close to expected values for primates of their size. The accelerometers recorded at 40 Hz, collecting >6.9 million records per axis for each individual. Individual DBA metrics were calculated and regressed against DLW daily energy measures. From this regression, we found a consistent and significant linear relationship (R^2^ = 0.46) between DLW and DBA, which could be improved by the incorporation of activity and resting time information (R^2^ = 0.52). Our results provide the first estimates of total daily energy expenditure for black lion tamarins and a validation of the method for estimating energy expenditure through accelerometers. Given the similar levels of total energy expenditure of captive and wild primates, this method can now be used in the field to estimate the energy cost of black lion tamarin movements in its natural environment.

**1 Summary statement:** Dynamic body acceleration was validated against doubly labelled water in black lion tamarins, showing it is a useful tool for measuring free-living energy demands.

## 3 Introduction

Quantifying energy expenditure in wild animals is fundamentally important in ecological research, such as in movement and behavioural ecology (e.g., Jeanniard-du-Dot et al., 2017b; Mathot et al., 2019) and life history studies (e.g., Jørgensen et al., 2016), including to understand the mechanisms underlying population dynamics (e.g., Andersen and MacMahon, 1981; Wearmouth et al., 2012) or responses to environmental and climate change (e.g., Fuller et al., 2016; Norin and Metcalfe, 2019). However, quantifying energy expenditure of wild animals is very challenging using standard methods, which rely either on invasive techniques (Butler et al., 2004) or on methods that are impractical in the field, such as those based on respirometry techniques (Pagano et al., 2018). A breakthrough was achieved by the doubly labelled water method (DLW; Speakman, 1997), which allows researchers to estimate average daily energy expenditure of free-living animals. Whilst the DLW method has been used on a wide variety of species (reviewed in McGrosky and Pontzer, 2023), it provides only a measure of average daily expenditure and does not allow the partition into the contributing parts at shorter time scales, such as different behavioural activities (e.g., running vs. walking), requiring other methods to explore energy expenditure between and within species (McGrosky and Pontzer, 2023).

Acceleration-based metrics such as Dynamic Body Acceleration (DBA), obtained from animal-borne accelerometers, are a powerful proxy for assessing fine-scale energy expenditure of free-living animals (reviewed in Wilson et al., 2020). This method was first proposed in a study with humans (Meijer et al., 1989; Montoye et al., 1983), and tested in animals two decades later, by calibrating the DBA-derived measures against metabolic rates using respirometry (Pagano et al., 2018; Wilson et al., 2006). Since then, DBA has been applied to predict costs of movement for a wide variety of species (Halsey et al., 2009; Wilson et al., 2020) – e.g., teleost fishes (Wright et al., 2014), cane toads (Halsey and White, 2010), cormorants (Laich et al., 2011; Stothart et al., 2016; Wilson et al., 2006), penguins (Hicks et al., 2020), hawks (Van Walsum et al., 2020), polar bears (Pagano and Williams, 2019), seals (Jeanniard-du-Dot et al., 2017a, 2017b), and canids (Tatler et al., 2021), but this is still lacking for important groups such as primates (McGrosky and Pontzer, 2023).

The total daily energy spent by an endothermic vertebrate (birds and mammals) is the sum of the energy expenditure used for the maintenance of bodily functions (basal metabolic rate; Mathot and Dingemanse, 2015), for digestion (specific dynamic action; Hawkins et al., 1997), for thermoregulation at temperatures below the lower critical limit of the thermoneutral zone (thermoregulatory thermogenesis; Scholander et al., 1950), for somatic production (such as growth and reproduction; Karasov, 1992), and for general patterns of activity, such as locomotion or movement (Christian et al., 1997) – see also review in McGrosky and Pontzer (2023). The costs of each of these components, especially those associated with basal metabolic rate, specific dynamic action and thermoregulation have been widely measured using the standard technique of indirect calorimetry, mostly by inferring heat production from measurements of gas exchange (Lighton, 2008). Although respirometry measurements have a long tradition of at least 200 years, this technique requires the confinement of the animal to a chamber or the use of oxygen masks, impractical for free-ranging animals, thus precluding energy quantification as the animals go about their daily routines.

The use of isotopes, such as DLW, circumvents some of the problems associated with respirometry (Carlson and Moreno, 1992). These techniques are based on the fact that animal bodies are in a constant flux of nutrients and resources, with substances being incorporated and eliminated at different rates. Thus, if the rate of elimination of a given substance is proportional to the metabolic rate, then this isotope technique allows for the measurement of the overall energy expenditure of free-ranging animals based on the rate of elimination of that substance (Bradshaw, 2003). Several isotopes have been used for this purpose, but the DLW technique that uses isotopes of oxygen (^18^O) and hydrogen (^2^H or ^3^H) is by far the most common method used to measure overall energy expenditure of free-ranging animals (Speakman, 1997). The DLW has however some key limitations that may hinder its application in wild animals, for being invasive and requiring recapturing of the same individuals after a very short period (often after 24 to 48 hours), and it can only provide measures of average daily energy expenditure.

Recently, acceleration-based metrics such as Dynamic Body Acceleration (DBA), obtained from animal-borne accelerometers, have been used to measure rates of energy expenditure of free-ranging animals (reviewed in Wilson et al., 2020). The reliability of this technique to trace rates of energy expenditure has been validated against standard respirometry methods (Enstipp et al., 2011, 2016; Ladds et al., 2017; Wright et al., 2014) and against the DLW method (Dalton et al., 2014; Elliott et al., 2012; Hicks et al., 2020; Jeanniard-du-Dot et al., 2017a, 2017b; Pagano and Williams, 2019; Stothart et al., 2016; Sutton et al., 2021). Although validation against these two methods attested for the reliability of the DBA as an indirect metric to assess total rates of energy expenditure, it should be noted that when validated against DLW, the reliability tends to be lower. This is because DLW measures the total energy spent by the animal during the monitoring period, while DBA measurements reflect only the costs associated with active movements, i.e., muscular activity, such as locomotion, movement and, perhaps, the work associated with shivering thermogenesis. In this sense, the DBA method is relevant for use in animals where the dynamic part (Wilson et al., 2020), i.e., movement, is a relevant component of the total daily energy spent, and where animal-attached devices are manageable in the field.

Here we aimed to validate the DBA method for estimating energy expenditure in primates, using black lion tamarins, *Leontopithecus chrysopygus* (Mikan, 1823), as our study species. Black lion tamarins are endangered small monkeys (body mass <800 g), that live in small family groups of two to eight individuals and move about in quadrupedal walking and leaps through the forest canopy as they forage for fruit and animal prey (Rezende et al., 2020). Due to their dependence on trees, a feature common to all Neotropical primates, they can be affected by changes in forest structure due to habitat fragmentation and degradation, such as occurring in the Brazilian Atlantic Forest, their native habitat (Ribeiro et al., 2009). Such habitat changes directly alter resource availability and distribution, which are mainly responsible for primate movement decisions (Trapanese et al., 2019). Thus, estimating movement costs can provide key information for the design of conservation actions, such as evaluating effects of forest fragmentation and planning forest restoration based on a forest structure that benefits tamarins’ movements.

Given that energy expenditure measured on free-living primates is comparable with those measured in primates in captivity (Pontzer et al., 2014), we were able to do a more controlled and less invasive experiment with captive tamarins. We used the standard procedure of combining the DLW method (Speakman and Racey, 1988; Speakman and Hambly, 2016) with the DBA method (reviewed in Wilson et al., 2020) for validation Once validated, we used the fine-scale information from the accelerometer data to determine the daily rhythm (day and night phases) and the fine-scale activity and resting patterns within each phase. Then, we used the latter to decompose the observed daily energy expenditure rates (based on dynamic acceleration) into the contributions of day/night phases and activity/resting behaviours in black lion tamarins to assess individual variability and improve the DLW/DBA energy rates relationship.

## 4 Materials and Methods

### 4.1 *Ex situ* Experiment with accelerometers and DLW

To validate the use of animal-attached sensor activity metrics to estimate energy expenditure in black lion tamarins, we conducted a study on captive animals at the Rio de Janeiro Primate Centre (CPRJ, Guapimirim, RJ, Brazil). Working in controlled environments with confined animals allows for more precise data collection and reduces the impact of multiple captures on wild animals.

In March 2018, ten healthy, adult black lion tamarins (eight males and two nonpregnant, non-lactating females, age 1.5 to 16 years old, weight 520–690 g; Table A1 in Supplementary information) were equipped for over 48 hours with “Daily Diaries” biologging units (DD; Wildbyte® Technologies Ltd., Swansea, UK) (Williams et al., 2020; Wilson et al., 2008). These units contain tri-axial accelerometer (providing information on posture and movement, and DBA measures) and magnetometer (providing orientation to magnetic north) sensors, and environmental temperature and pressure sensors (the latter providing information on altitudinal movements), recording at a range (40 – 4 Hz) of sub-second frequencies (Table 1). In particular, the accelerometer data provided the individual DBA metrics used as proxy for energy expenditure (Wilson et al., 2020).

**Table 1.** A list of variables the Daily Diary (DD) collected with corresponding recording frequency, units’ measurement, and range.

| **Channel** | **Recording frequency (Hz)** | **Measurement** |
| --- | --- | --- |
| Accelerometer X axis Sway (Side – Side) | 40 | -16 to 16 *g* |
| Accelerometer Y axis Surge (Forward – Backward) | 40 | -16 to 16 *g* |
| Accelerometer Z axis Heave (Upward – Downward) | 40 | -16 to 16 *g* |
| Magnetometer axis 1 | 16 | Max. of earth's magnetic field |
| Magnetometer axis 2 | 16 | Max. of earth's magnetic field |
| Magnetometer axis 3 | 16 | Max. of earth's magnetic field |
| Barometric pressure | 4 | 100 to 2000 mbar |
| External Temperature | 4 | -20 to 60°C |

The device was attached to the animal using a backpack-style harness made of nylon cord holding a plastic housing (Figure 1) containing the DD circuit board powered by a 3.7V 123 mAh rechargeable lithium battery. The shape and size of the devices used were developed specifically for lion tamarins, considering its weight, ergonomics, and anatomy restrictions (Figure 1), and based on models currently in use by field projects with similar species (Savage et al., 1993). Each tag was colour-coded externally to allow visual individual identification during the experiment (Figure 1). The total weight of the backpack was 18 g, making up 2.6–3.5% of the animals’ body mass (500–700 g) and staying within the ‘5% rule’ recommended by the ethical guidelines for mammals (Sikes and Gannon, 2011; Wilson et al., 1996).

**Figure 1.**
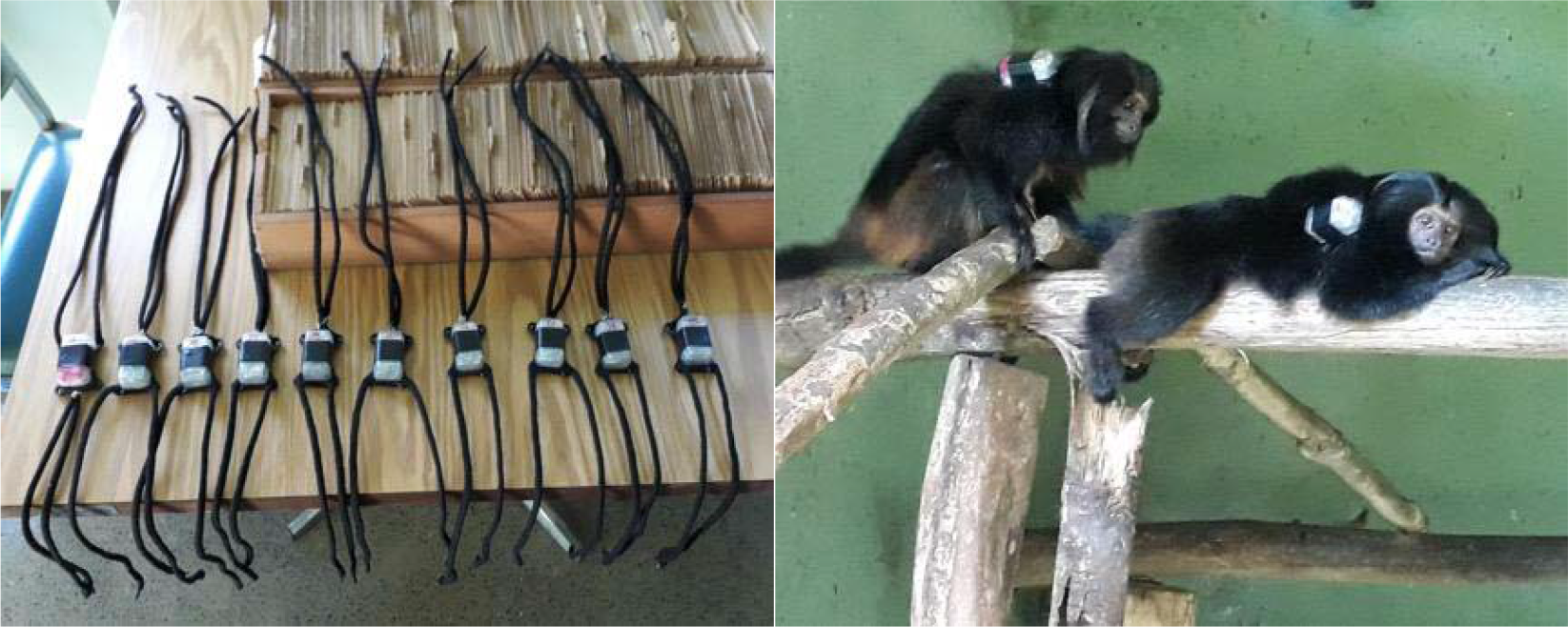
Daily Diaries used on the black lion tamarins at Rio de Janeiro Primate Centre (CPRJ) (left). Two black lion tamarins with the daily diaries on their backs, at CPRJ, Guapimirim, RJ, Brazil (right).

Immediately before attaching the DD, each individual was weighted (Pesola® 1000 g, d = 10 g) and subsequently injected intraperitoneally with 0.4 to 0.53 ml (depending on body mass) of a water solution enriched with 65% of ^18^O and 35% of ^2^H, which was weighed to 4 d.p. Based on the enrichment level of the amount of water injected and the body mass (Speakman 1997), a 2-3-hour period was allowed for the injected solution to equilibrate with the body water pool. After this period, a 100ul blood sample was taken from the femoral vein into capillary tubes, which were flame sealed and stored at 2-4°C before being sent to analysis. Soon after this initial blood sampling and anaesthesia recovery, individuals were returned to their respective enclosures (3.0 x 6.0 x 2.5 m), where they were free to move and develop their usual behaviours. Two days later, individuals were removed from their outdoor cages, had their body mass measured again, and a second blood sample was taken and processed in a similar way as the first bleeding procedure. All animals were anaesthetised during the whole procedure to reduce handling stress, with a combination of ketamine 10 mg/kg (Cetamin 10%, Syntec) and midazolam 1 mg/kg (Midazolam 5 mg/mL, Hipolabor) intramuscularly (Carpenter and Marion, 2017).

The blood samples were analysed in the DLW Resource Centre, at the Zoology Department of the University of Aberdeen (Scotland), following Westerterp (2017) to estimate the elimination rates of ^18^O (ko) and ^2^H (kd). Analysis of the isotopic enrichment of blood was performed blind, using a Liquid Isotope Water Analyser (Los Gatos Research, USA) (Berman et al 2013). Initially the blood encapsulated in capillaries was vacuum distilled (Nagy 1983), and the resulting distillate was used for analysis. Samples were run alongside five lab standards for each isotope and International Standards to correct for daily variations. A single-pool model was used to calculate rates of CO_2_ production as recommended for use in animals less than 5 kg in body mass (Speakman 1993). The isotopic dilution space was estimated from both ^2^H and ^18^O, and is equivalent to the total body water pool, expressed as a percentage of the body mass prior to injection. Using these data, the amount of CO_2_ produced (VCO_2_) was calculated and averaged on a daily basis (mlCO_2_/day). Since the initial blood sampling was taken right after the equilibrium period, the Plateau method was used to calculate VCO_2_, following equation 7.17 of Speakman (1997; see also Speakman and Król, 2005). We also tested the Intercept method and used three different equations for each method to get daily energy expenditure (DEE) values (Estimate 1 = equation 35 *sensu* Lifson and McClintock, 1966; Estimate 9 = equation 7.17 *sensu* Speakman, 1997; and Estimate 4 = equation A6 *sensu* Schoeller et al., 1986, modified by Schoeller, 1988; Table A1 in Supplementary information), which provided very similar and highly correlated estimates (Figure A1 in Supplementary information). Thus, for further analysis we used the results obtained from Estimate 9 Plateau, which is considered the most appropriate estimate for small mammals. Finally, a respiratory quotient of 0.8 was used to calculate the total DEE (in kJ/day) from VCO_2_ (Speakman 1997). This value of DEE was then later compared with the data from the DD to validate the use of DBA as a measurement of energy expenditure.

### 4.2 Accelerometer data processing and DBA estimation

Data management, visualisation, and analysis of the Daily Diary biologging data was done using the specialised software Daily Diary Multiple Trace Graphing Tool (DDMT; Wildbyte® Technologies Ltd., Swansea, UK). The programme produces interpolated time-based plots to show how acceleration and other channels change over time at high resolution and allows to quickly subset the data, calculates DBA and other derived metrics, and identifies different behavioural phases using the Boolean time-based decision-tree template method (Wilson et al., 2018).

For the analyses, we considered the data collected whenever the animal was free to move on its own in the enclosure, but not including the time when the animal was being held or manipulated, as this would otherwise directly influence the accelerometer records. This interval coincided with the same time interval between the two blood samples taken for the DLW estimate, assuring full comparability. From the acceleration data, an activity metric called Vectoral Dynamic Body Acceleration (VeDBA) was calculated using DDMT via the following equation:

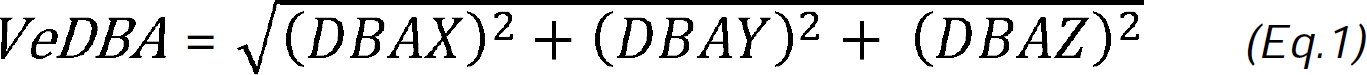

where DBA is the dynamic body acceleration in three dimensions: X or sway (side to side), Y or surge (back and forth) and Z or heave (up and down). Dynamic acceleration is calculated for each record by subtracting the static from the raw acceleration, to obtain a measure that reflects only active movements, using a 2-second moving average window (i.e., 80 records at 40 Hz), centred on each record, to estimate the static acceleration signal (Wilson et al., 2020).

To reduce the noise in the metric, we applied a running mean across 200 data points (5 seconds) to produce smoothed VeDBA (sVeDBA). Other DBA metrics derived from the acceleration data, such as Overall Dynamic Body Acceleration (ODBA) and smoothed ODBA, were also used (Wilson et al., 2020) but, since these metrics were highly correlated (see Figure A1 in Supplementary information), we focused our analyses on sVeDBA.

### 4.3 Daily Rhythm Phases: the distinction between day and night phases and active and inactive behaviours

Further analysis on the accelerometer data was carried out to identify active or inactive phases (Figure 2). As the sensor is very sensitive to even small movements of the animal, reflected in higher values of *g*, differences in the signal between inactivity and activity are generally very clear (Shepard et al., 2008), being at very low *g* values during inactivity and markedly larger and usually more variable when the animal is active (Figure 2).

**Figure 2.**
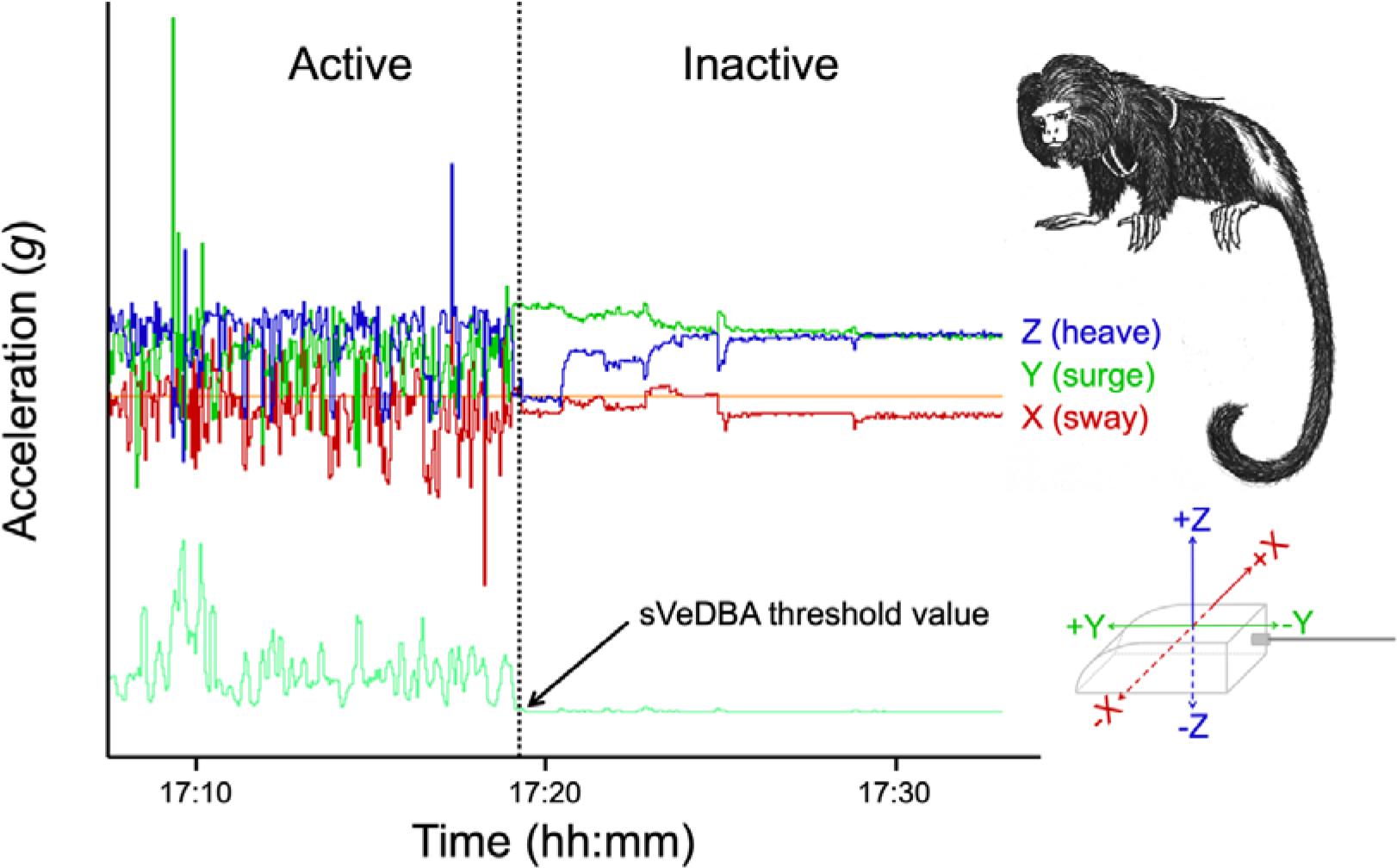
Patterns observed on the accelerometer vectors (red, green, and blue lines) and smoothed Vectorial Dynamic Body Acceleration (sVeDBA; cyan line) when the black lion tamarins (top-right) are active or inactive. The vertical dotted line indicates the exact time where the transition to the inactivity (night) phase happened, which coincides with sVeDBA values below the threshold chosen here to identify resting behaviour (indicated by the arrow). The orange line corresponds to acceleration = 0. The three axes shown on the device (bottom-right) indicate the respective vectors when the tamarin is in this horizontal position. Tamarin illustration: Gustavo Libardi.

To determine the daily rhythm of each individual and identify activity (day) and inactivity (night) cycles, we visually inspected the data to find start and end of activity hours based on a threshold value of sVeDBA. To define this threshold value that would represent a change between activity and inactivity (Figure 2), initially we exported the sVeDBA values for each record over the monitoring period and subdivided this data into day and night phases based on the civil twilight times for that day in Guapimirim, Rio de Janeiro (05h36 to 18h19). Then we run descriptive statistics (mean, median, and quartiles) for each dataset corresponding to the night phase for the ten black lion tamarins. Based on the third quartile values of sVeDBA, we defined the threshold value that would correspond to the highest value to describe inactivity (sVeDBA < 0.04).

Using DDMT, we marked the data points based on this threshold value, with all data points with a sVeDBA value lower than the threshold considered as ‘inactive’, and higher as ‘active’. From the length of these bouts, we could clearly identify the transition times between active and inactive phases as whenever the activity records were longer than the inactivity ones. The opposite criterion was used for the start of the inactive phase. Based on these times, we subdivided the data into these two phases of the daily rhythm, which here were named as “DAY” and “NIGHT” phases, respectively. This way we could estimate the length of each phase of the daily rhythm for each individual, and the proportion of time that they spent on each day/night phase.

In addition to the simple rule to separate the active daytime and the inactive nighttime, we defined two supplementary rules to identify fine-scale changes between active and inactive behaviours within each of the day and night time phases (e.g., short bouts of posture change during sleep, or resting bouts during the day), using time-based behaviour identification rules (Wilson et al., 2018). To clearly decipher between inactive resting periods (rather than pauses) during the DAY, the animal had to be defined as inactive by the sVeDBA threshold (<0.04) for a minimum interval of a minute (2400 records) – listed as Rule 1, or the rule that detected the daytime resting behaviour. This was then repeated to ensure the species was active during the NIGHT, also avoiding flinching and other instantaneous active behaviours, using the same threshold value to define activity (sVeDBA > 0.04 for over a minute) – named Rule 2, or the rule that detected the night time activity behaviour. On DDMT, we marked all records following these rules, applying Rule 1 for the DAY phase (1-marked records are resting, 0-marked are active) and Rule 2 for the NIGHT phase (0-marked records are resting, 1-marked are active).

After marking the behaviours, we could create six subsets of data that would include all (T), resting (R), and activity (A) events, in each phase of the daily rhythm (DAY vs. NIGHT): T_DAY_, R_DAY_, A_DAY_, T_NIGHT_, R_NIGHT_, and A_NIGHT_, respectively.

### 4.4 Energy Expenditure estimates

To calculate DBA-based daily energy estimates, we summed the sVeDBA values for each record during the length of the experiment (∼48 h), generating a total sVeDBA, and estimated the daily sVeDBA proportionally to a 24-hour period. This is the same approach hence, over the same time period, as used to obtain the Daily Energy Expenditure estimates from the DLW data. These daily sVeDBA values for each tamarin were regressed against the DLW daily energy measures (DEE) to validate the former (“stats” package, R Core Team, 2022) in R 4.0.2 (R Core Team, 2020). Standard residual diagnostic checks were used, significance of regression slopes was evaluated using standard methods for linear models (Crawley et al., 2012) and fit was evaluated using the R^2^ metrics (Elliott et al., 2012; Stothart et al., 2016). As sVeDBA values are highly correlated to the other DBA metrics, i.e., VeDBA, ODBA and sODBA (Wilson et al., 2020) (see Figure A1 in Supplementary information), we chose to use sVeDBA (*sensu* Stothart et al., 2016) for validating the DD energy expenditure estimates against the DLW ones.

We also checked the impact that collecting accelerometer data at a different frequency would have on the regression using 20Hz datasets (to allow comparison and conversion in case other devices collect data at different frequencies). For this, we subsampled the dataset collected at 40Hz, taking only the even or odd rows to simulate a dataset collected at 20Hz. Again, we regressed these DBA values against the DLW daily energy measures.

Subsequently, to improve the relationship between DBA and DLW estimates, we quantified the contribution of activity and resting behaviours to the daily energy spent by the black lion tamarins. For this, we estimated total and daily sVeDBA, and associated time lengths, for each day/night phase and active/inactive behaviour. We summed the sVeDBA values of the records within each subset and adjusted it proportionally to the time length of each. Based on these sVeDBA values partitioned in different behaviours and their respective time length, similarly to the method by Jeanniard-du-Dot et al. (2017a), we estimated behaviour-specific energy expenditure. For this, we fit the following model for all our tamarins to obtain the parameter estimates for each behaviour (a, b, c and d):

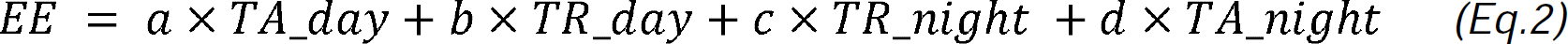

where:

*EE* = total energy spent during the experiment

*a, b, c, d* = rate of energy expenditure for each behaviour

*T_i_* = Time spent in each behaviour

We also ran alternative models including as covariate the different combinations of the proportion of time spent in each behaviour and the phases of the daily rhythm. We used AIC-based multi-model comparison methods, and the best fitted model was selected based on the Akaike Information Criterion (AIC). Models with ΔAIC < 2 were considered equally plausible (Burnham and Anderson, 2002).

All analyses were undertaken using the software DDMT, R 4.0.2 (R Core Team, 2020) and R Studio 1.3.1073 (RStudio Team, 2020).

## 5 Results

### 5.1 Validation of DBA metrics

The results from the DLW blood samples analysis provided a mean Daily Energy Expenditure measure of 353.5 kJ/day (SD=73.8 kJ/day, or 84.5 ± 17.6 kcal/day), with the lowest values obtained from the two females (Table 2). We did not observe a direct association between body mass and variation in energy expenditure (R^2^= 0.27, p > 0.05). The analysis of the Daily Diaries data provided a mean daily sVeDBA of 279,706.2 *g* (SD=58,747.4 *g*), with no differences associated with the sex or weight of the individuals (Table 2). We found a consistent and significant linear relationship between sVeDBA and DEE values (F1,8 = 6.739, R^2^ = 0.46, p = 0.03; Figure 3). DEE values (in kJ/day) increased by 0.00085 kJ for each *g* of DBA, according to the following equation:

**Figure 3.**
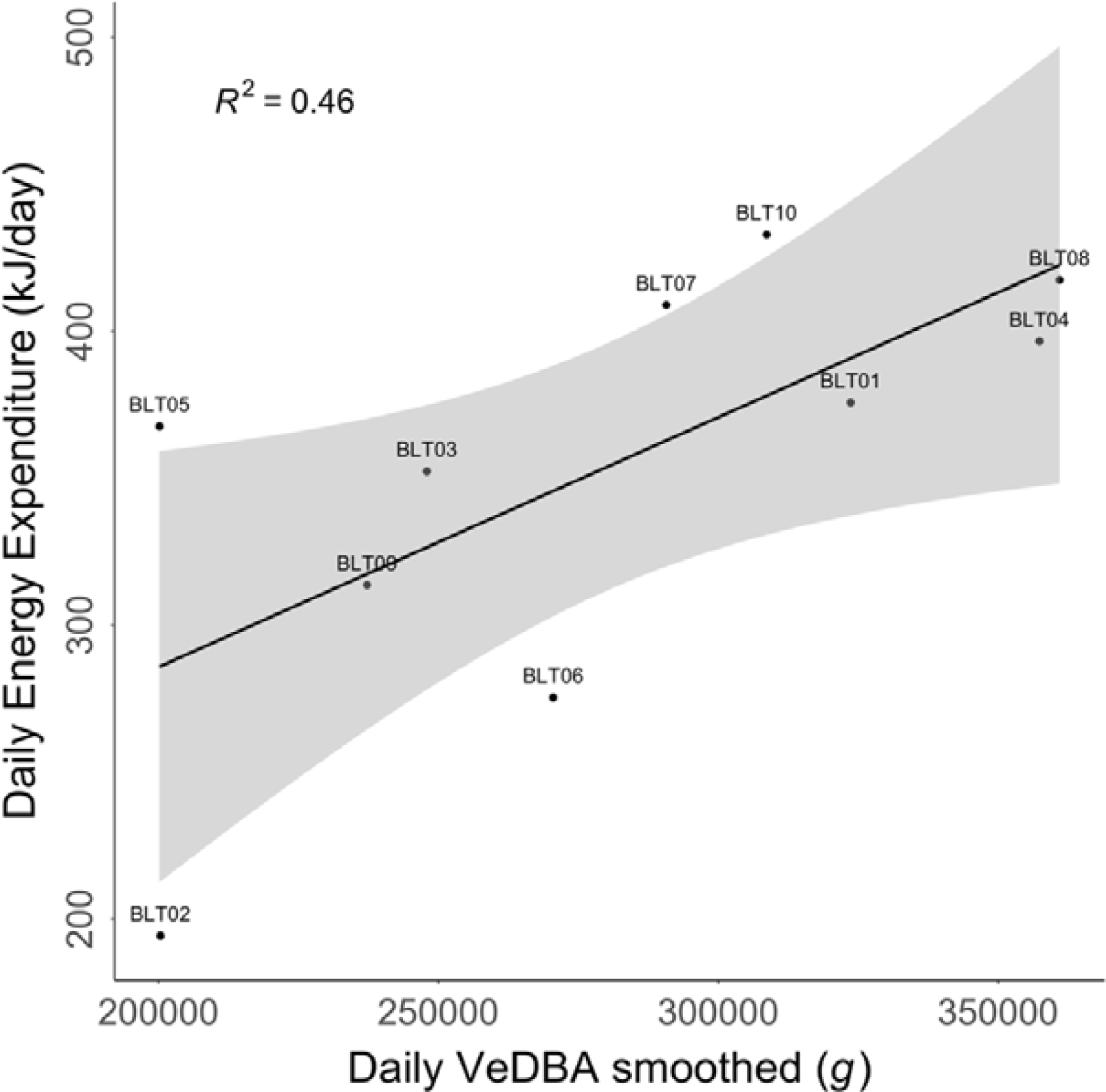
Linear regression between daily energy expenditure (DEE), in kJ per day, and smoothed Vectorial Dynamic Body Acceleration (sVeDBA), in g (acceleration unit) per day. The ID of each individual is shown next to each point on the graph. The grey area indicates the 95% C.I.

**Table 2.** Daily energy expenditure (DEE) in kJ and kcal per day and Daily smoothed Vectorial Dynamic Body Acceleration (sVeDBA) in acceleration units (g) for each black lion tamarin (ID). The values of mean body mass (BM) in kilograms (kg), consider the tamarins’ weights at the beginning and at the end of the experiment. The mean and standard deviation (SD) values for each variable, considering all animals of the experiment (All), the females (F) and the males (M), are presented on the last three rows, respectively.

| ID | Sex | Mean BM (kg) | DEE (kJ/day) | DEE (kcal/day) | Daily sVeDBA (g) |
| --- | --- | --- | --- | --- | --- |
| BLT01 | M | 0.5225 | 375.7 | 89.8 | 324,023.50 |
| BLT02 | F | 0.5125 | 194.3 | 46.4 | 200,697.50 |
| BLT03 | M | 0.6725 | 352.3 | 84.2 | 248,294.60 |
| BLT04 | M | 0.6075 | 396.6 | 94.8 | 357,749.60 |
| BLT05 | M | 0.6525 | 367.6 | 87.9 | 200,531.50 |
| BLT06 | F | 0.5325 | 275.3 | 65.8 | 270,877.30 |
| BLT07 | M | 0.5950 | 408.9 | 97.7 | 291,076.10 |
| BLT08 | M | 0.6000 | 417.4 | 99.8 | 361,334.90 |
| BLT09 | M | 0.5975 | 313.6 | 75.0 | 237,620.60 |
| BLT10 | M | 0.6000 | 432.9 | 103.5 | 309,007.80 |
| MEAN<br>±SD | All | 0.5893 ±0.0529 | 353.5 ±73.8 | 84.5 ±17.6 | 280,121.3 ±58,737.7 |
|  | F | 0.5225 ±0.0141 | 234.8 ±57.3 | 56.1 ±13.7 | 235,787.4 ±49,624.6 |
|  | M | 0.6059 ±0.0444 | 383.1 ±38.9 | 91.6 ± 9.3 | 291,204.8 ±58,155.8 |

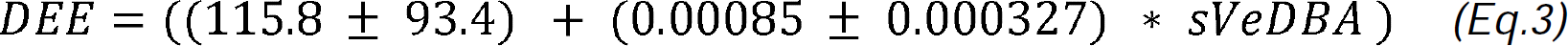

When we rerun the regression using 20-Hz subsampled datasets, the results showed a similar relationship, with the same intercept value (115.8) but a proportional change on the slope, where DEE values increase by 0.0017 kJ for each *g* of DBA. We obtained the same results for either odd or even-rows datasets (see Figure A2 in Supplementary information).

### 5.2 Behaviour-specific Energy Expenditure

The black lion tamarins were monitored for a mean period of 47h10m12s (±26m07s) (Table 3). During this period, they spent, on average, 48.2% (±2.8%) of the time on the DAY phase and 51.8% (±2.8%) on the NIGHT phase of the daily rhythm (Table 3). The animal that spent the longest time on the NIGHT phase (BLT02) was a 15-year-old female that was most affected by the anaesthetic process and was kept in a controlled environment for recovery until the next morning.

**Table 3.** Length of the activity (DAY) and inactivity (NIGHT) phases, proportion of time spent in each phase, and total monitoring period (time of the device on the animal) over the two days of the experiment. The horizontal lines on the table group the animals that were kept in the same enclosures. The mean (±SD) length and proportions are presented on the last row. The shortest and longest lengths and monitoring periods are highlighted (bold).

| ID | Length of<br>NIGHT phase<br>(hh:mm:ss) | Length of<br>DAY phase<br>(hh:mm:ss) | % Time on<br>NIGHT<br>phase | % Time on<br>DAY<br>phase | Total<br>monitoring<br>period |
| --- | --- | --- | --- | --- | --- |
| BLT01 | 24:55:29 | 22:43:36 | 52.3% | 47.7% | 47:39:05 |
| BLT02 | **27:49:20** | **19:11:38** | **59.2%** | **40.8%** | 47:00:58 |
| BLT03 | **23:21:00** | 23:37:44 | **49.7%** | **50.3%** | 46:58:44 |
| BLT04 | 23:29:15 | 23:20:15 | 50.2% | 49.8% | 46:49:30 |
| BLT05 | 24:37:55 | 22:03:10 | 52.8% | 47.2% | **46:41:05** |
| BLT06 | 24:04:08 | 23:44:09 | 50.3% | 49.7% | **47:48:17** |
| BLT07 | 23:52:21 | **23:55:09** | 50.0% | 50.0% | 47:47:30 |
| BLT08 | 23:34:55 | 23:43:54 | 49.8% | 50.2% | 47:18:49 |
| BLT09 | 24:12:38 | 22:36:49 | 51.7% | 48.3% | 46:49:27 |
| BLT10 | 24:15:39 | 22:32:54 | 51.8% | 48.2% | 46:48:33 |
| **MEAN** | 24:25:16 | 22:44:56 | 51.8% | 48.2% | 47:10:12 |
| **(±SD)** | ±01:17:41 | ±01:23:56 | ±2.8 | ±2.8 | ±00:26:07 |

Despite the similar time spent in each phase of the daily rhythm, the mean sVeDBA for the DAY was four times higher (T_DAY_ = 78.7% ±4.7; T_NIGHT_ = 21.3% ±4.7), with most of this sVeDBA corresponding to active behaviour (76.0% ±5.5). We also observed that there is activity during the NIGHT, although very infrequent both in time (0.7% ±0.5) and in contribution to total sVeDBA (1.5% ±1.6), while resting during the DAY accounted for 8.1% (±4) of the time and 2.6% (±1.6) of the sVeDBA (Figure 4). The same female previously mentioned (BLT02) presented the lowest sVeDBA values for A_DAY_ (see Table A2 and A3 in Supplementary information).

**Figure 4.**
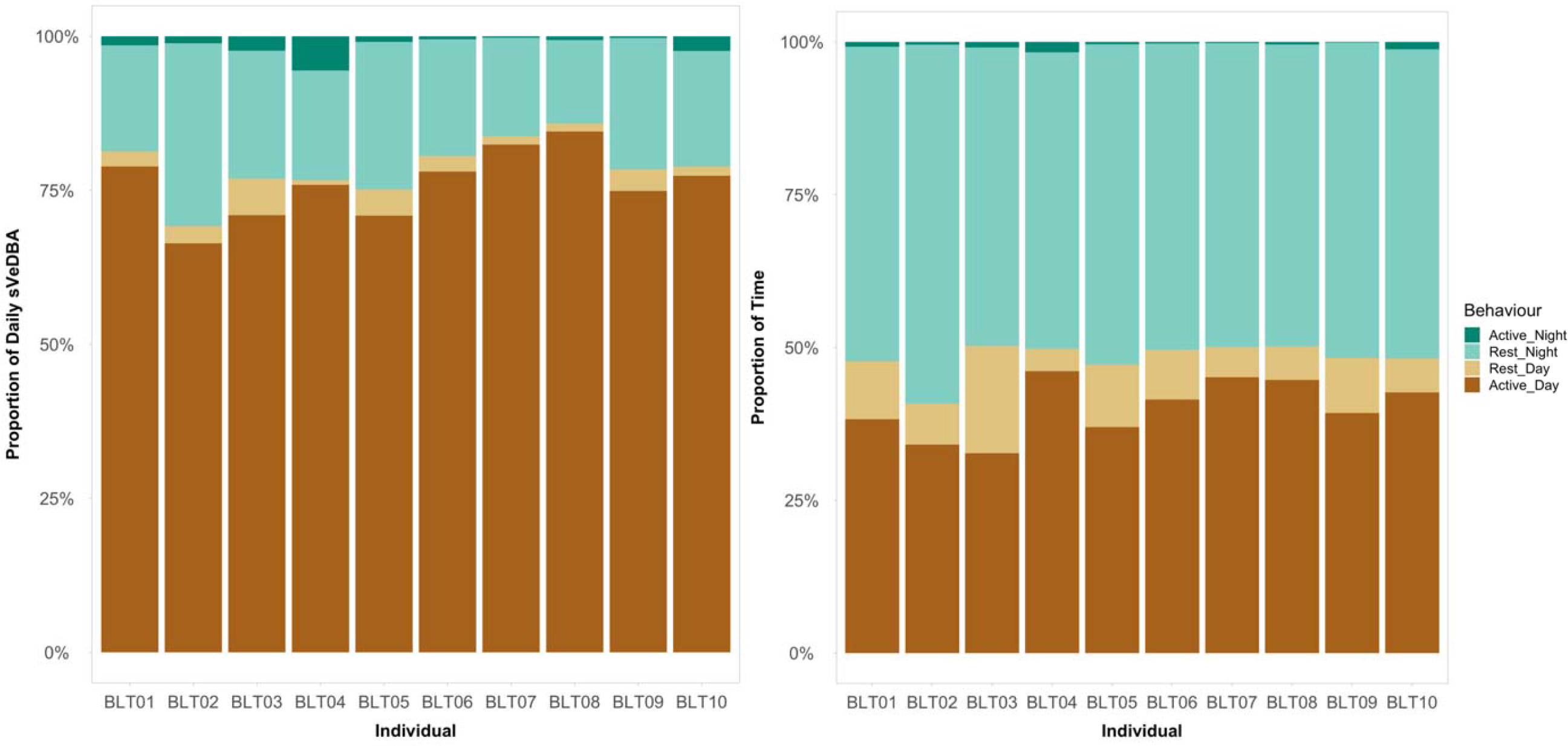
Proportion of daily sVeDBA (left) and time (right) spent in each behaviour (active or resting, during the day or night) within 24 hours for each black lion tamarin.

We fit the model for Equation 2 using Energy Expenditure (EE) values based on DLW metrics for the total monitoring period. We found a negative parameter estimate for the resting behaviour at night, but the confidence interval included zero (Table 4). As the p-value associated with this behaviour and phase of the daily rhythm were higher than 0.05, we ran alternative models including as covariate the different combinations of the proportion of time spent in each behaviour and the phases of the daily rhythm (Table 4). We found that activity and the day phase, alone or combined, were the main factors that influenced energy expenditure rates in black lion tamarins (p ≤ 0.01). The model that included only the total proportion of time in activity, independent of the phase (m5), was the best predictor of EE (Table 4). This model did not have a better fit than the one from Equation 2 (m1; F-test p > 0.3) but presented a much lower AIC value (ΔAIC = 14.3; Table 4).

**Table 4.** Parameter estimates of the models on the relationship between energy expenditure (in kJ) and the costs of activity and resting behaviours, during day and night, for black lion tamarins. Shaded row corresponds to the best-fit model according to AIC.

| Model | Dependent variable | Parameter<br>(in days) | Estimate<br>(in kJ/day) | SE | p | R<sup>2</sup> | AICc | ΔAIC |
| --- | --- | --- | --- | --- | --- | --- | --- | --- |
| m0 | TEE (DLW) | totTime | 353.13 | 52.55 | **<0.001** | 0.958 | 132.69 | 4.79 |
| m1 | TEE (DLW) | timeActDay | 1289.3 | 866.01 | **0.011** | 0.977 | 142.20 | 14.30 |
|  |  | timeRestDay | 868.83 | 1291.37 | 0.151 |  |  |  |
|  |  | timeRestNight | -500.23 | 775.7 | 0.166 |  |  |  |
|  |  | timeActNight | 3206.96 | 9020.86 | 0.418 |  |  |  |
| m2 | TEE (DLW) | timeActDay | 814.65 | 178.56 | **<0.001** | 0.973 | 131.38 | 3.48 |
|  |  | timeActNight | 4052.55 | 8981.41 | 0.329 |  |  |  |
| m3 | TEE (DLW) | totTimeDay | 1302.35 | 785.08 | **0.005** | 0.976 | 130.16 | 2.26 |
|  |  | totTimeNight | -532.05 | 732.3 | 0.132 |  |  |  |
| m4 | TEE (DLW) | totTimeAct | 917.15 | 585.57 | **0.007** | 0.971 | 132.14 | 4.24 |
|  |  | totTimeRest | -36.03 | 405.37 | 0.843 |  |  |  |
| m5 | TEE (DLW) | totTimeAct | 866.00 | 44.51 | **<0.001** | 0.974 | **127.90** | **0.00** |

Legend: EE = total Energy Expenditure on the monitoring period; SE = Standard Error; p = significance level; R<sup>2</sup> = adjusted R<sup>2</sup>; AICc = Akaike Information Criterion corrected for small sample size; ΔAIC = difference in AIC score between the best model and the model being compared; totTime = total monitoring time; timeRestDay/Night = time spent resting during the respective phase; timeActDay/Night = time spent in activity during the respective phase; totTimeDay/Night = total time spent on the respective phase; totTimeRest/Act = total time spent on the respective behaviour.

According to the selected model (m5), we calculated the energy spent in activity (in kJ) by multiplying the respective parameter estimate from the model (in kJ/day) by the respective time spent active (in days) for each black lion tamarin (i.e., EE_ACT_ (kJ) = 866 * totTimeAct). Then, we regressed this value against the sVeDBA value for this behaviour (A_DAY_ + A_NIGHT_; Table S2 in Supplementary information) and found that this regression greatly improved our VeDBA/EE relationship (R^2^ = 0.69, p = 0.003; Figure 5), when compared to the original one (R^2^ = 0.46, p = 0.03; see Figure 3).

**Figure 5.**
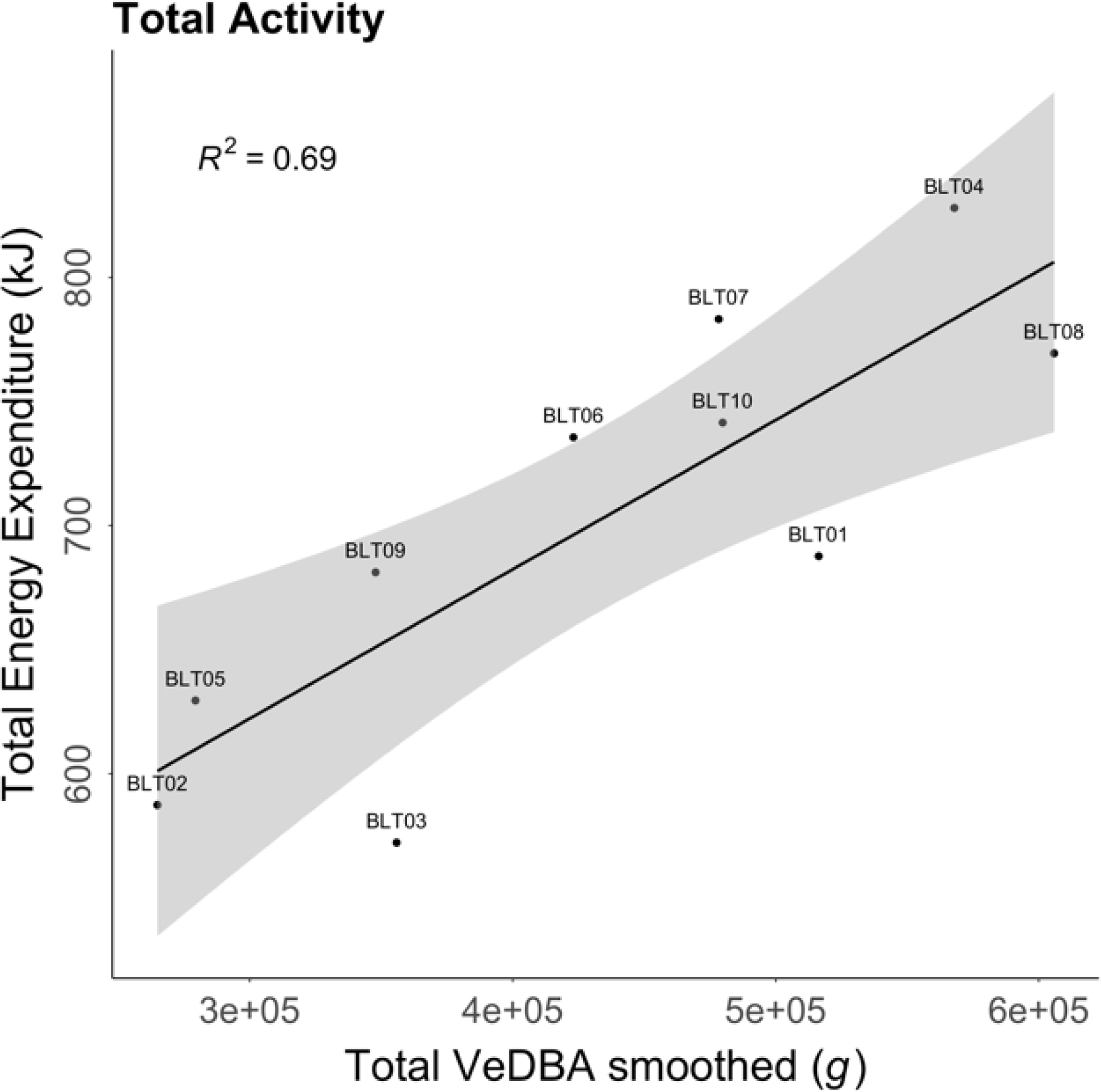
Linear regression between energy expenditure (EE), in kJ, and smoothed Vectorial Dynamic Body Acceleration (sVeDBA), in g, including only activity records. The ID of each individual is shown next to each point on the graph. The grey area indicates the 95% C.I.

Finally, from this relationship, we derived equation 4 to predict total energy expended (TEE) by the animals based on the sVeDBA values from activity, and equation 5 to predict daily energy values (DEE):

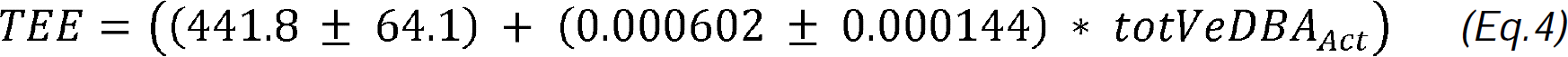

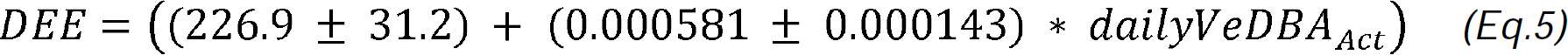

When we compared the predicted values from equations 4 and 5, with the measured ones using the DLW method, we found that the relationship improved from the original one (respectively, R^2^ = 0.52 and R^2^ = 0.5, p = 0.02).

## 6 Discussion

We provide the first estimates of metabolic rate for captive black lion tamarins. Values obtained from DLW are in close agreement with expected values from the literature (84.6 kcal ± 5.2, range: 50.2 kcal ± 3.2 – 142.6 kcal ± 8.6; Table A4 in Supplementary information; Pontzer et al., 2014), including known parameters from closely related marmoset species. They are also similar to the values estimated for wild golden lion tamarins, using the same method (87.9 kcal ± 10.8; A. P. Cruz-Neto, unpublished data). When we regressed the DLW results against the DBA values, we found a consistent linear and significant relationship between the two measures. The accuracy we found is within the range of values reported in the literature for DLW/DBA relations, with lower R^2^ compared to other studies in penguins (0.72 in Hicks et al., 2020), cormorants (0.81 in Stothart et al., 2016) and polar bears (0.70 in Pagano and Williams, 2019), but higher R^2^ when compared to wild fur seals (0.36 in Jeanniard-du-Dot et al., 2017b), and similar to those observed for semi-captive sea lions (0.47 in Fahlman et al., 2008).

When we partitioned the sVeDBA values into activity and resting, we saw that nearly 80% of it is associated with active states, reinforcing the assumption that black lion tamarin movement is a major component of the total daily energy spent. Particularly, energy expenditure (expressed in DBA metrics) related to activity during the night accounted for 1.5% of total sVeDBA. Very low activity rate during night is expected for the species as callitrichids are known to reach states of torpor during sleep, probably as an energy-saving mechanism (Thompson et al., 1994). It is worth noting that we were not able to accurately estimate resting metabolic rates (RMR) from our methods. Thompson et al. (1994) reported, for captive golden lion tamarins, resting-phase RMR equivalent to 2.46 kcal.h^-1^.kg^-1^ (43.4 ±7.4 kcal.day^-1^, mean weight: 0.733 kg ± 0.099). Considering physiological similarities between these species, if we apply this same rate to black lion tamarins, we find an average RMR of 34.8 kcal.day^-1^ (± 3.1). This is nearly 40% of the total DEE for males and over 54% for females. The latter agrees to another study with wild female golden lion tamarins that presented RMR estimations that were more than half of their DEE rates (Miller et al., 2006). More precise estimations of RMR for each phase of the daily rhythm, and their relative contribution to the DEE in tamarins can be achieved if data on ambient temperature is included in the analysis (Thompson et al., 1994).

Both females of our study presented the lowest DEE values, a common fact in primates, when it comes to non-lactating females (Key and Ross, 1999). Differences in DEE rates between males and females are more prominent in primates with sexual dimorphism, especially due to body size differences, but still present in similar-sized different-sex individuals, which is the case for black lion tamarins (Coelho, 1974; Key and Ross, 1999). However, our low sample size does not allow us to state that this difference was significant.

The results of our models reinforced that activity was the best predictor of energy expenditure, while resting energy expenditure rates were neither accurate nor significant, presenting negative estimates and high standard errors. This reflects that the DBA method is usually not appropriate for estimating energy expenditure rates of behaviours that account for subtle or insignificant movement, even if collecting data at a very fine scale (Gleiss et al., 2011; Green et al., 2009; Jeanniard-du-Dot et al., 2017). Following the approach taken by Jeanniard-du-Dot and collaborators (2017a), we could increase the accuracy of our relation by using only the VeDBA values related to activity, excluding resting, and pairing with the time each animal spent active within the monitoring period. Accelerometer data allow to infer fine scale behaviour of tagged animals to a resolution that goes well beyond the difference between moving *versus* resting used here, by defining algorithms derived from acceleration-sensor metrics representative of these behaviours (Wilson et al., 2018). It was already demonstrated that slopes of VeDBA/EE relationships can vary by activity type (Hicks et al., 2020; Jeanniard-du-Dot et al., 2017a). Therefore, additional analyses of the accelerometer data could partition black lion tamarin activity into more specific behaviours, especially the more costly ones, such as walking and travelling, but also grooming, jumping, feeding, etc., to define more specific activity budgets and further refine the use of DBA metrics for estimating energy expenditure in wild tamarins.

The validation of this methodology for estimating the energy expenditure of black lion tamarins based on DBA-metrics sets a precedent for the use of accelerometers in studies of this nature in small primates in different contexts, whether in captivity or in the wild. The fine-scale decomposition of daily energy expenditure into contributions of day/night and active/resting bouts highlights the unique potential to use the DBA-metrics to compare energy expenditure of populations living in forests with different degrees of fragmentation and forest structure, which is vital to provide a more quantitative evidence base for conservation and mitigation measures. It will be particularly interesting to assess if DEE of individuals of different populations changes linearly or non-linearly (e.g., if there are sudden breakpoints) with forest fragmentation and if this is further affected by seasonal changes, effects of group size etc., further improving the evidence base for more efficient management and conservation measures for black lion tamarins in human-dominated landscapes.

## 7 List of symbols and abbreviations

AIC: Akaike Information Criterion
BLT: black lion tamarin
BM: body mass
DBA: Dynamic Body Acceleration
DD: Daily Diary (device)
DDMT: Daily Diary Multiple Trace Graphing Tool (software)
DEE: Daily Energy Expenditure
DLW: Doubly Labelled Water
EE: Energy Expenditure
ODBA: Overall Dynamic Body Acceleration
RMR: Resting Metabolic Rate
sODBA: smoothed Overall Dynamic Body Acceleration
sVeDBA: smoothed Vectorial Dynamic Body Acceleration
TEE: Total Energy Expenditure
VeDBA: Vectorial Dynamic Body Acceleration

## 8 Acknowledgements

We thank Phil Hopkins (Swansea University) for helping in designing and building the tag housings and attachment, and Mark Holton for helping to prepare the Daily Diary loggers. We also thank Gustavo Simões Libardi, who made the black lion tamarin illustration included in Figure 3.

## 10 Author contributions

Gabriela C. Rezende, Ariovaldo P. Cruz-Neto, Luca Börger, Rory Wilson and Laurence Culot contributed to the study’s conception and design. The daily diary loggers, housings, attachments, and individual marks where prepared by Luca Börger; all DLW method procedures were prepared by Ariovaldo P. Cruz-Neto. Data collection was performed by Ariovaldo P. Cruz-Neto, Luca Börger, Guilherme S. T. Garbino, Alcides Pissinatti and Silvia Bahadian. Gabriela C. Rezende, Luca Börger, Ariovaldo P. Cruz-Neto, James Redcliffe, Catherine Hambly and John Speakman developed the data analysis. The first draft of the manuscript was written by Gabriela C. Rezende, Luca Börger and Ariovaldo P. Cruz-Neto, and all authors commented on previous versions. All authors read and approved the final manuscript.

## 12 Data Availability

All data supporting the analyses will be made available on FigShare at the time of publication.

